# A widespread methylotroph acyl-homoserine lactone synthase produces an atypical quorum sensing signal

**DOI:** 10.1101/2022.10.31.514548

**Authors:** Dale A. Cummings, Mike Wallace, Andrew G. Roberts, Aaron W. Puri

## Abstract

Pink pigmented facultative methylotrophs of the genera *Methylorubrum* and *Methylobacterium* are omnipresent bacteria often found associated with plants. Despite their widespread occurrence, the molecular details of how these organisms interact with each other and their environment remain understudied. We analyzed genes encoding *N*-acylhomoserine lactone quorum sensing signal synthases in published genomes of these bacteria and determined that the product of the largest group of signal synthases had not been characterized. We subsequently identified this *N*-acylhomoserine lactone product using inverse stable isotopic labeling (InverSIL), which revealed an atypical signal. We then demonstrate in one representative strain that this signal activates its cognate LuxR-family transcription factor and is produced in a positive feedback loop. These results reveal a previously undescribed yet widespread signal used by pink pigmented facultative methylotrophs, which helps us understand the chemical ecology of these important bacteria.

---

The bacterial genera *Methylobacterium* and *Methylorubrum* primarily consist of pink pigmented facultative methylotrophs (PPFMs) that are widespread in nature and the built environment.^1,2^ These bacteria are often associated with plants, both in the phyllosphere^3^ and as endophytes,^4^ and have been shown to have important roles in plant growth promotion.^5,6^ Bacteria from these genera have also been identified as the causative agents of some hospital-acquired infections.^7^ Despite this recognized importance, the molecular details of how PPFMs interact with each other and their environment are understudied.

Many Gram-negative proteobacteria including PPFMs use *N*-acylhomoserine lactones (acyl-HSLs) as quorum sensing (QS) signals.^8–11^ Acyl-HSL signals are produced by LuxI-family synthases and exhibit variation in their acyl chain length and substituents but maintain a common HSL core originating from methionine.^12,13^ Acyl-HSLs are bound by LuxR-family transcription factors that subsequently regulate gene expression to coordinate group behaviors,^10^ including biofilm formation^14^ and antibiotic production.^15^ Characterizing these QS systems can help us understand the molecular details of PPFM interactions.

We recently reported the use of an inverse stable isotopic labeling (InverSIL) approach to identify methylotroph acyl-HSL QS signals.^16^ In this approach, we grow a bacterial strain on a ^13^C-labeled version of its carbon source to generate a fully ^13^C-labeled culture. We can then feed this culture ^12^C-precursors and detect their incorporation into natural products by mass spectrometry, thereby eliminating problems with ^13^C-labeled precursor availability. In the case of methylotroph acyl-HSLs, bacteria are grown on ^13^C-methanol and then fed ^12^C-methionine, which is incorporated into the HSL portion of these signals.

In this work, we identified a widespread acyl-HSL synthase gene in publicly available genomes of the genera *Methylobacterium* and *Methylorubrum*. We then used InverSIL to identify the acyl-HSL product of this synthase family and confirmed this result in multiple acyl-HSL synthase representatives from different strains. When we isolated and determined the structure of this compound, it revealed a previously undescribed signal. In one representative strain we then demonstrate that this acyl-HSL signal activates its cognate LuxR-family transcription factor and is produced in a positive feedback loop. These findings provide new information about the chemical ecology of these ubiquitous bacteria.

## A widespread methylotroph acyl-HSL synthase produces an uncharacterized product

To determine the diversity of signals produced by PPFMs, we organized the 271 *Methylobacterium*/*Methylorubrum* LuxI-family acyl-HSL synthases in the Joint Genome Institute’s Integrated Microbial Genomes and Microbiomes (IMG/M) system^17^ using sequence similarity networking,^18,19^ as has been done previously with acyl-HSL synthases.^20^ We used an alignment score threshold of 85, which produces isofunctional groups of acyl-HSL synthases (Amy Schaefer, University of Washington, personal communication) **(Fig. 1)**. In other words, all members of each synthase cluster are predicted to produce the same acyl-HSL signal. While some clusters contain synthases that have been characterized (MlaI, MsaI1-3)^8,16^ the largest cluster does not have a characterized representative, and therefore a widely conserved signal in this group of organisms cannot be predicted.

**Figure 1.**
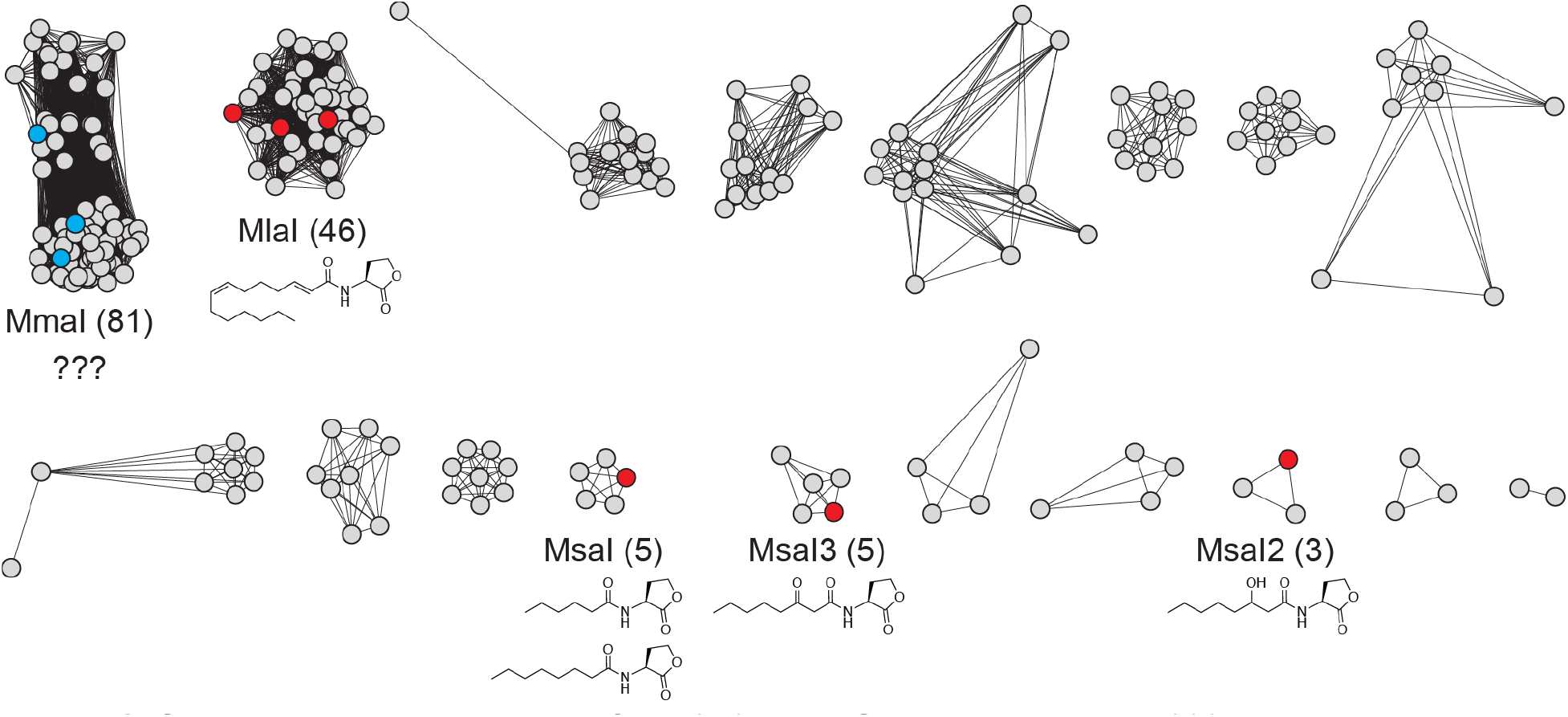
Sequence similarity network of the 271 acyl-HSL synthases in the 200 *Methylobacterium* and *Methylorubrum* genomes in the IMG/M system. Red nodes show acyl-HSL synthases with previously characterized products. The structure of the major product is shown when reported. Cyan nodes show acyl-HSL synthases characterized in this work. The number of nodes in a cluster are indicated in parentheses. 21 singletons are not shown.

We next wanted to identify the acyl-HSL product of this cluster of synthases. PPFM strains often contain multiple annotated *luxI*-family synthase genes in their genomes,^9,21^ which can make it difficult to link production of an acyl-HSL to a specific synthase. *Methylobacterium* sp. strain 88A^22^ possesses only one synthase and it is found in the largest cluster, enabling us to unambiguously link this synthase with its product.

We used InverSIL to identify the acyl-HSL signal produced by 88A. When we grew 88A on ^13^C-methanol, we identified a feature with an *m/z* of 314 that decreased four *m/z* units when we added ^12^C-methionine, which is consistent with incorporation of four carbon atoms from methionine into an HSL **(Fig. 2A)**. In the ^12^C-methanol control this feature had an *m/z* of 298, indicating this feature likely corresponds to a protonated acyl-HSL containing 16 carbons total, with 12 present in the acyl chain.

**Figure 2.**
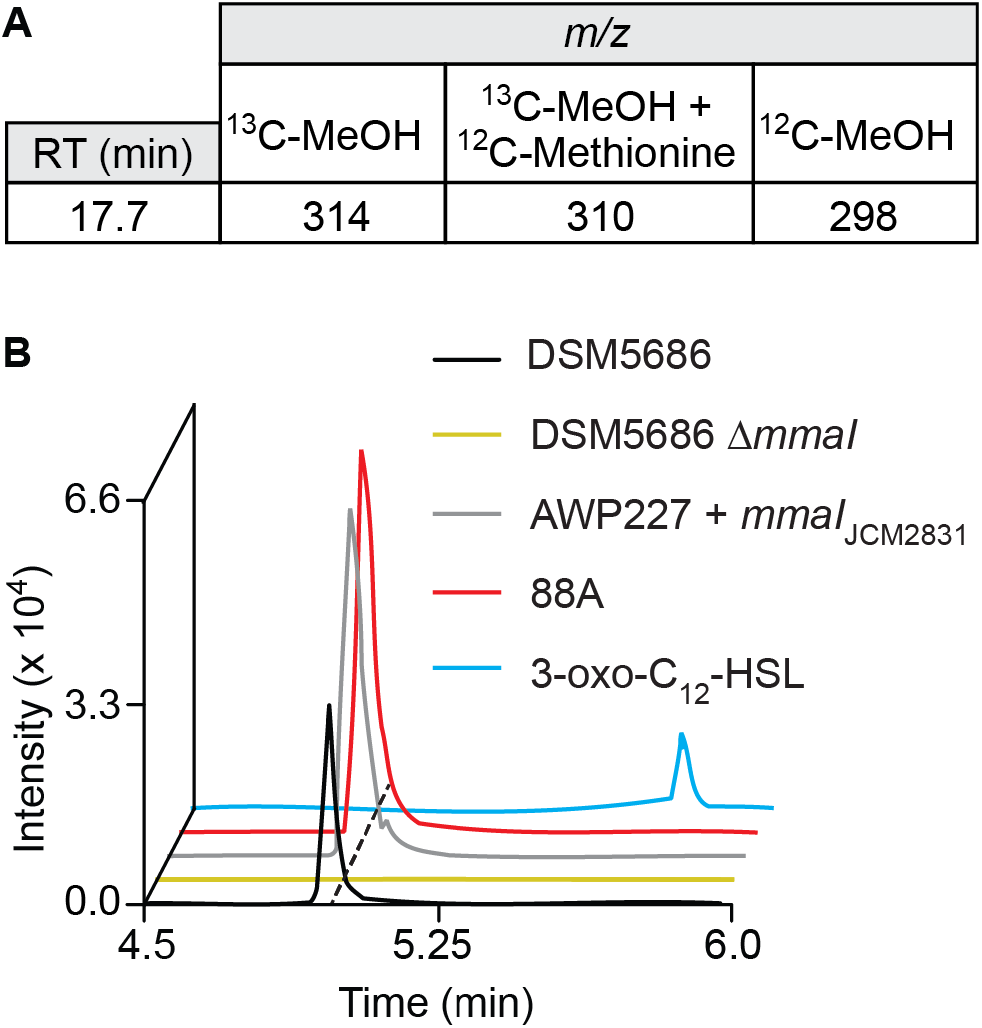
(A) Inverse labeling of 88A culture identified a feature that incorporates methionine. (B) Extracted ion chromatogram of supernatant extracts for the listed strains for *m/z* 298.2013, corresponding to protonated 3*R*-OH-5*Z*-C_12:1_-HSL. Mass tolerance < 5 ppm.

To verify this result, we also examined the products of other representatives of this acyl-HSL synthase cluster using high resolution LC-MS/MS. When we heterologously expressed the synthase representative from *Methylobacterium radiotolerans* JCM2831^23^ (Mrad2831_5763) in the strain AWP227, where the endogenous QS genes *mlaR* and *mlaI* have been deleted,^16^ we observed an identical feature **(Fig. 2B)** with the same fragmentation pattern as the feature from 88A **(Table S1)**. We also observed an identical feature in a culture of *Methylobacterium fujisawaense* DSM5686,^24^ which also contains a representative synthase from this cluster (Ga0373204_3345), and this feature was not present in a deletion mutant of this synthase **(Fig. 2B)**. Together, these results indicate that this feature is the product of the widespread cluster of acyl-HSL synthases **(Fig. 1)**.

The observed high-resolution *m/z* of the feature is consistent with the known signal *N*-(3-oxododecanoyl)-*L*-homoserine lactone (3-oxo-C_12_-HSL) (observed [M+H]^+^ = 298.2013, expected [M+H]^+^ = 298.2013, 0 ppm). However, a 3-oxo-C_12_-HSL standard exhibited a longer retention time **(Fig. 2B)**, indicating that the target acyl-HSL signal product is likely a constitutional isomer of 3-oxo-C_12_-HSL that to our knowledge had not been previously described. We therefore decided to isolate and structurally characterize this compound.

### Characterization of the acyl-HSL product of the major cluster reveals an atypical structure

We chose DSM5686 to scale up production of the target acyl-HSL. We grew 30 one-liter cultures and extracted the supernatant with acidified ethyl acetate. The dried extract was then separated by C_18_ solid phase extraction and purified by HPLC guided by the compound mass. This method yielded 1.2 mg of the pure signal. To increase the signal-to-noise ratio for structural elucidation using NMR spectroscopy, we also grew three one-liter cultures of DSM5686 using ^13^C-methanol as the sole carbon source. Structural elucidation was performed by ^1^H, ^13^C, and various 2D NMR spectroscopy experiments (**Figs. 3** and **S1-S6**,**Table S2**). This analysis revealed a hydroxyl group on carbon 3 and an olefin between carbons 5 and 6 of the aliphatic chain. The ^3^J_HH_-coupling of 8.9 Hz enabled us to assign the olefin in a *cis* configuration.

**Figure 3.**
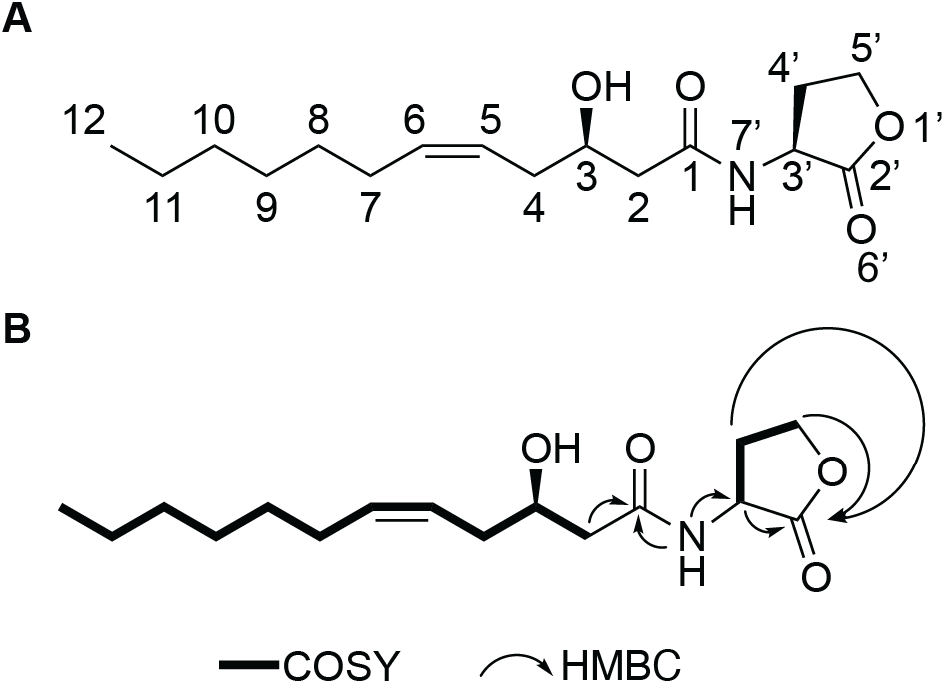
(A) Structure of 3*R*-OH-5*Z*-C_12:1_-HSL (B) 2D NMR structural assignments.

We next determined the absolute stereochemistry of this compound. Marfey’s analysis revealed an *L*-HSL **(Table S3)**, which is typical for acyl-HSL QS signals.^25^ To determine the stereochemistry of the secondary alcohol substituent, we first reduced the olefin using palladium on carbon in the presence of H_2_ gas to create 3-OH-C_12_-HSL, which we then subjected to methanolysis to produce methyl 3-hydroxydodecanoate **(Fig. S7)**.^26^ Using chiral gas chromatography, we compared this methyl ester to commercially available racemic methyl 3-hydroxydodecanoate as well as methyl 3*R*-hydroxydodecanoate **(2)**, which we synthesized via an enantioselective reduction of methyl 3-oxododecanoate **(Fig. S8)**.^27^ This analysis showed the hydroxyl in the *R* configuration **(Fig. S9)**, which has also been noted for other acyl-HSL signals where stereochemistry has been determined.^26,28^ Together, this demonstrates that the acyl-HSL product of this cluster of acyl-HSL synthases is *N*-(3*R*-hydroxy-5-*cis*-dodecenoyl)-*L*-homoserine lactone (3*R*-OH-5*Z*-C_12:1_-HSL). We therefore name the cluster of synthases that produces this acyl-HSL signal MmaI, for *<u>M</u>ethylobacterium*/*<u>M</u>ethylorubrum* <u>m</u>edium-chain <u>a</u>cyl-HSL.

### 3*R*-OH-5*Z*-C_12:1_-HSL activates the transcription factor MmaR

To determine if the signal is bioactive, we constructed an acyl-HSL QS reporter assay.^29^ The *mmaI* gene is co-located in the DSM5686 genome with a gene encoding a LuxR-family transcription factor (Ga0373204_3344), which we term *mmaR*. We inserted *mmaR* into the genome of our previously constructed heterologous expression strain AWP227 downstream of the native *mlaR* promoter leftover in that strain. We then conjugated a plasmid into this strain containing the red fluorescent reporter gene *mScarlet* driven by the DSM5686 *mmaI* upstream region. LuxI-family acyl-HSL synthases are often positively autoregulated by their cognate LuxR-family transcription factors,^11,29^ and therefore we hypothesized mScarlet would be expressed when the signal is added to this reporter strain.

Upon signal addition to the reporter strain, we observed an increase in red fluorescence with an EC_50_ of 0.2 nM **(Fig. 4)**. This is over one thousand-fold lower than the 720 nM ±10 nM we quantified in early stationary phase cultures of DSM5686. We also determined that this signal is produced when we grow DSM5686 on methanol but not the multicarbon substrate succinate, as has been previously observed for acyl-HSL QS signals produced by MlaI in *Methylorubrum extorquens* AM1 **(Fig. S10)**.^8^

**Figure 4.**
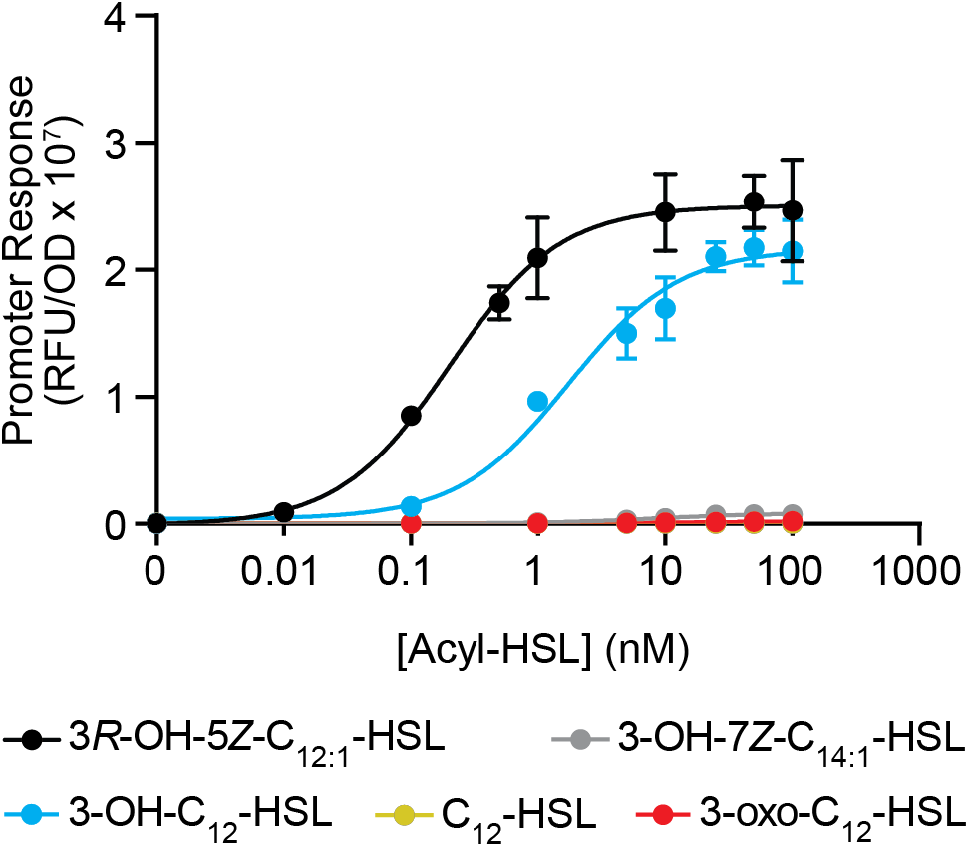
Activation of transcription factor MmaR by acyl-HSL signals. RFU: Relative fluorescence units. OD: Optical density at 600 nm. Data show the mean and standard deviation of five cultures and are representative of two independent experiments.

When we tested four structurally related, commercially available acyl-HSLs (3-oxo-C_12_-HSL, 3-OH-C_12_-HSL, C_12_-HSL, and 3-OH-7Z-C_14:1_-HSL) in the reporter strain, only 3-OH-C_12_-HSL activated MmaR at a concentration similar to the native signal (EC_50_ 1.7 nM) **(Fig. 4)**. Notably, the 3-OH-C_12_-HSL signal we tested was a mixture of four diastereomers (*N*-(3*RS*-hydroxydodecanoyl)-*DL*-homoserine lactone), so it is possible that a single diastereomer may have an increased potency. Together, these results demonstrate that *3R*-OH-5*Z*-C_12:1_-HSL can activate the transcription factor MmaR and that this signal is produced in a positive feedback loop. This strategy of inserting *luxR*-family transcription factor genes into AWP227 and then testing various acyl-HSL signals and plasmid-contained receptor binding sites is also generalizable and can be used to characterize more LuxR-family transcription factors from alphaproteobacteria in the future.

## Summary and Conclusion

PPFMs are widespread, metabolically versatile bacteria with reported interactions with plants, other bacteria, and humans. Here we identified and characterized a QS system that is widely conserved among PPFMs of the genera *Methylobacterium* and *Methylorubrum*. We determined that this QS system produces and responds to the signal 3*R*-OH-5*Z*-C_12:1_-HSL, which to our knowledge has not been previously described. Because this signal is a constitutional isomer of the more commonly described signal 3-oxo-C_12_-HSL, our work highlights the importance of rigorous characterization of acyl-HSL QS signals by LC-MS/MS in the absence of full NMR spectroscopic studies. Furthermore, the discovery of new acyl-HSL signaling systems can help synthetic biologists use exogenous signals to encode orthogonal regulatory circuits in organisms.^30^ Our analysis of the QS systems in *Methylobacterium* and *Methylorubrum* also shows there are many other signals to be characterized **(Fig. 1)**, some of which may also have new structures.

## ACKNOWLEDGEMENTS

This work was supported by National Institutes of Health grants R00 GM118762 and R35 GM147018 (to A.W.P.). We thank members of the Puri Lab for reading the manuscript and for helpful discussions. We thank Peter Flynn (University of Utah) for helpful discussions about NMR structural elucidation. We thank Emily Mevers (Virginia Tech) for advice on determining the hydroxyl stereochemistry of *3R*-OH-*5Z*-C_12:1_-HSL and Stefan Schulz (TU Braunschweig) for advice on acyl-HSL methanolysis. We thank Dakota Brady (University of Utah) for help with initial development of the acyl-HSL Marfey’s analysis procedure in our lab. The *mScarlet* PCR template pMRE-Tn7-135 was a gift from Mitja Remus-Emsermann (Addgene plasmid #118553).^31^

## AUTHOR CONTRIBUTIONS

A.W.P., A.G.R., and D.A.C. and M.W. designed the experiments. D.A.C. and M.W. performed the experiments. A.W.P. and D.A.C. wrote the manuscript. All authors edited and approved of the final version of the manuscript.

## METHODS

See the Supporting Information.

