## SupplementaryInformation for "A widespread methylotroph acyl-homoserine lactone synthase produces an atypical quorum sensing signal"

### METHODS

**Key reagents.** $^{13}C$-labeled methanol was purchased from Cambridge Isotope Laboratories. 3-OH-$C_{12}$-HSL was purchased from Millipore Sigma. All other acyl-HSLs were purchased from Cayman Chemical.

**Routine bacterial culturing.** Strains used in this study are listed in **Table S4**. *Escherichia coli* strains were grown in lysogeny broth (LB) at 37 ºC. *Methylorubrum extorquens* AWP227 derivatives, *Methylobacterium* sp. strain 88A, and *Methylobacterium fujisawaense* DSM5686 were grown at 30 ºC in modified ammonium mineral salts (AMS) medium,[1] with the addition of 0.01% (w/v) yeast extract for 88A and DSM5686. Modified AMS contains 0.2 g $L^{-1}$ $MgSO_4 \cdot 7H_2O$, 0.2 g $L^{-1}$ $CaCl_2 \cdot 6H_2O$, 0.5 g $L^{-1}$ $NH_4Cl$, 30 µM $LaCl_3$, and 1X trace elements. 500X trace elements contains 1.0 g $L^{-1}$ $Na_2$-EDTA, 2.0 g $L^{-1}$ $FeSO_4 \cdot 7H_2O$, 0.8 g $L^{-1}$ $ZnSO_4 \cdot 7H_2O$, 0.03 g $L^{-1}$ $MnCl_2 \cdot 4H_2O$, 0.03 g $L^{-1}$ $H_3BO_3$, 0.2 g $L^{-1}$ $CoCl_2 \cdot 6H_2O$, 0.6 g $L^{-1}$ $CuCl_2 \cdot 2H_2O$, 0.02 g $L^{-1}$ $NiCl_2 \cdot 6H_2O$, and 0.05 g $L^{-1}$ $Na_2MoO \cdot 2H_2O$. A final concentration of 4 mM phosphate buffer pH 6.8 and 50 mM $^{12}C$- or $^{13}C$-methanol were added prior to use and cultures were shaken at 200 rpm.

**Acyl-HSL synthase sequence similarity networking.** On January 27, 2022 all genomes of the genera "*Methylobacterium*" and "*Methylorubrum*" were downloaded from the IMG/M system, returning 200 genomes. These genomes were then searched for genes containing pfam00765, returning 271 genes. Amino acid sequences were exported in FASTA format and analyzed using the EFI workflow using an alignment score threshold of 85. The network was visualized without collapsing nodes containing 100% amino acid identity with Cytoscape 3.9.1 using the Prefuse Force Directed OpenCL Layout by alignment score.

**Inverse stable isotopic labeling (InverSIL) experiments.** Inverse labeling and subsequent analysis was performed as previously described.[2] Exponentially growing bacterial cultures were pelleted at 16,100 rcf for one minute and resuspended in growth medium with no carbon source. Subsequently, three separate six milliliter cultures were inoculated with the resuspended strain at a starting OD of 0.02. The $^{12}C$-carbon source was added to one culture, the $^{13}C$-carbon source to the second, and the $^{13}C$-carbon source plus 500 nM $^{12}C$-methionine to the last culture. The carbon sources used were 50 mM methanol, 50 mM methylamine, or 50% (v/v) methane. Cultures were grown until reaching stationary phase (OD of approximately 0.8) and then were centrifuged at 4,800 rcf for ten minutes. The resulting supernatant was extracted twice with an equal volume of ethyl acetate containing 0.01% (v/v) acetic acid, and the combined organic extract was evaporated to dryness using a nitrogen stream and stored at -20 ºC until analysis by LC-MS.

**LC-MS for acyl-HSL signal detection.** Dried culture supernatant extracts were resuspended in 200 microliters of 1:1 water:acetonitrile, and subsequently 65 microliters were injected onto an Agilent 1260 Infinity liquid chromatography system connected to an Agilent 6120 single quadrupole mass spectrometer operating with positive polarity and a mass range of 150-1500 *m/z*. A Waters Xselect HSS T3 column (2.5 µm particle size, 2.1 mm x 50 mm) held at 30 ºC was used for reverse phase separation with a flow rate of 0.4 mL $min^{-1}$. Solvent A: Water + 0.1 % formic acid, Solvent B: Acetonitrile + 0.1% formic acid. Gradient: 0-2 min, 0% B. 2-32 min, 0-100% B. 32-35 min, 100% B. 35-36 min, 100-0% B. 36-38 min, 0% B.

**Inverse stable isotopic labeling (InverSIL) analysis.** Inverse labeling analysis was performed as previously described.[2] Raw data files in netCDF format were exported using Agilent OpenLab CDS (rev C.01.07). Features were detected using MZmine version 2.53[3] using the following workflow: 1. Mass detection (centroid, noise level 1.0E3). 2. ADAP chromatogram builder[4] (minimum group size 5 scans, group intensity threshold 1.0E3, min highest intensity 5.0E3), *m/z* tolerance 0.3). 3. Chromatogram deconvolution (local minimum search, chromatogram threshold 30%, search minimum 0.1 min, minimum relative height 10%, minimum absolute height 6.0E3, minimum ratio of peak top/edge 2, peak duration 0-2 min). 4. Adduct search (RT tolerance 0.1 min, adducts $[M+Na]^+$ and $[M+NH_4]^+$ selected, *m/z* tolerance 0.2, max relative peak height 200%). 5. Feature list rows filter (remove identified adducts). Subsequently, isotopes were removed from $^{12}C$ samples using the Isotopic peaks grouper (*m/z* tolerance 0.2, retention time tolerance 0.1 min, monotonic shape required, maximum charge 3, representative isotope most intense), and the three feature lists were aligned in the order $^{13}C$-carbon source + $^{12}C$-methionine, $^{12}C$-carbon source, $^{13}C$-carbon source using the Join aligner (*m/z* tolerance 0.3, weight of *m/z* 50, retention time tolerance 0.1 min, weight for retention time 50). The alignment was exported in .csv format with the row retention time as a common element and peak *m/z* as the data file element. Features containing the desired four *m/z* unit difference in the $^{13}C$-carbon source and $^{13}C$-carbon source + $^{12}C$-methionine samples were then detected using a custom Python script (available at https://github.com/purilab/inverse).

**Plasmid construction.** Plasmids used in this study are listed in **Table S5**. Primers used in this study are listed in **Table S6**. All plasmids were constructed using Gibson Assembly[5] and selection was performed with kanamycin (50 µg $mL^{-1}$).

**Genetic manipulation.** All gene locus tags in this manuscript refer to the Joint Genome Institute Integrated Microbial Genomes & Microbiomes data management system (JGI IMG/M).[6] Genetic manipulation of strain AWP227 was performed at 30 ºC. Sequence verified plasmids were conjugated into these strains using the *E. coli* donor S17-1.[7] 500 µL of exponentially growing cultures (OD 0.4-0.6) of the donor and recipient strains were pelleted at 16,100 rcf for one minute and resuspended in 500 µL sterile ultrapure $H_2O$. These strains were then pelleted again, and the two pellets were combined in a total volume of 50 µL sterile ultrapure $H_2O$. Next, the entire mixture was spotted onto an AMS agar plate containing 50 mM methanol and 10% (v/v) nutrient broth and incubated for two days. Successful conjugants were selected on AMS plates containing kanamycin (50 µg $mL^{-1}$). To construct the unmarked insertion mutant AWP348, kanamycin-resistant integrants (single crossovers) were restreaked and then plated on an AMS plate containing 50 mM methanol and 1% (m/v) sucrose for counterselection. The resulting colonies were screened for double crossovers by kanamycin sensitivity and colony PCR before the final mutant was verified by Sanger sequencing.

**High-resolution tandem mass spectrometry.** Mass spectrometry data were collected using a Waters Acquity I-class ultra-high pressure liquid chromatography (UPLC) instrument coupled to a Waters Xevo G2-S quadrupole time-of-flight mass spectrometer. An Acquity UPLC BEH C18 column (2.1 x 50 mm) was used for separation and resolving samples. Solvent A: Water + 0.1 % (v/v) formic acid, Solvent B: Acetonitrile + 0.1% (v/v) formic acid. The sample was eluted from the column using a ten minute linear solvent gradient: 0-0.1 min, 1% B; 0.1 - 10 min, 1-100% B. The solvent flow rate was 0.45 mL $min^{-1}$. Mass spectra were collected in positive ion mode, with following parameters: 3 kV capillary voltage; 25 V sampling cone voltage; 150 °C source temperature; 500 °C desolvation temperature; nitrogen desolvation at 800 L/hr. The fragmentation spectra were collected using the same parameters with a 10-25 eV collision energy ramp. The lockspray solution was 200 pg/µL leucine enkephalin. The lockspray flow rate was 6 µL/min. Sodium formate was used to calibrate the mass spectrometer.

**Large scale production and purification of 3*R*-OH-5*Z*-$C_{12:1}$-HSL.** 6 mL of an exponentially growing culture of *Methylobacterium fujisawaense* DSM5686 was used to inoculate 1 L AMS + 50 mM MeOH with a starting optical density of 0.02. A total of six 1 L cultures were grown at a time. These cultures were incubated at 30℃ and shaken at 200 rpm to stationary phase (approximately 72 hours). The culture was centrifuged at 4700 rpm for 10 min followed by extraction of the clarified supernatant with an equal volume of ethyl acetate acidified with 0.01% v/v acetic acid. The organic phase was then dried by rotary evaporation. Dried crude supernatant extracts were resuspended in 1 mL of 1:3 water:acetonitrile (ACN) and applied onto a preequilibrated Discovery $C_{18}$ solid-phase extraction (SPE) column (3 mL, 500 mg) then washed and eluted sequentially with 6 mL of 25%, 50%, 75%, and 100% ACN in water. The acyl-HSL was found in the 75% ACN fraction by LCMS. This fraction was dried and resuspended in 500 uL of 25% ACN in water and separated using an Agilent 1260 Infinity liquid chromatography system. A Waters SunFire C18 column (3.5 µm particle size, 46 mm x 100 mm) was used for reverse phase separation with a flow rate of 1 mL $min^{-1}$. Solvent A: Water + 0.1 % trifluoroacetic acid, Solvent B: Acetonitrile + 0.1% trifluoroacetic acid. Gradient: 0-1 min, 10-35% B. 1-21 min, 35-55% B. 21-22 min, 55-100% B. 22-30 min, 100% B. 30-31 min, 100-10% B. 31-36 min, 10% B.

***3R*-OH-*5Z*-$C_{12:1}$-HSL**: HRESIMS: $[M+H]^+$= 298.2013 (calculated for $C_{16}H_{28}NO_4^+$: 298.2013; 0.0 ppm). MS2: see **Table S1**. $^1H$ NMR (800 MHz, $CDCl_3$) and $^{13}C$ NMR (200 MHz, $CDCl_3$): see **Table S2**.

**Marfey's analysis of 3*R*-OH-5*Z*-$C_{12:1}$-HSL.** 3*R*-OH-5*Z*-$C_{12:1}$-HSL (0.1mg) was suspended in 250 µL of 6M HCl in water overnight at 110 °C to cleave the amide bond. This reaction was then lyophilized and the resulting solid was resuspended in 250 µL of saturated sodium bicarbonate. 16 µL of 1% (w/v) of Marfey's reagent (Sigma-Aldrich, 71478) was added and the mixture was allowed to react for 1 hour at 40 °C. The reaction was then quenched with 20 µL of 2M HCl in water and analyzed by LC-MS, where an *m/z* of 372 was found, corresponding to the $[M+H]^+$ of the hydrolyzed HSL with a Marfey's adduct. Retention times were compared to Marfey's derivatized *L*- and *R*-HSL standards.

**Catalyst preparation for synthesis of 3*R*-hydroxydodecanoic acid methyl ester.** The catalyst, (R)-$[RuCl_2(BINAP)]_2 \cdot NEt_3$, was prepared by a procedure adopted from Taber and Silverberg.[12] All steps were performed under inert atmosphere. Briefly, 40 mg Dichloro(1,5-cyclooctadiene) ruthenium(II) polymer (Sigma-Aldrich, 337331) and 100 mg (R)-BINAP (Sigma-Aldrich 295187) were dissolved in toluene (5 mL) and triethylamine (0.3 mL) was added. The mixture was refluxed (oil bath at 140 °C) for 4 hours, after which the solvent was removed under vacuum. The resulting red-orange powder was dissolved in 1 mL THF and immediately used in the next reaction.

**Reduction of methyl 3-oxododecanoate to 3*R*-hydroxydodecanoic acid methyl ester (2).** Methyl 3-oxododecanoate **(1)** (100mg) (TRC, M324720), methanol (0.5 mL) and the above catalyst (0.1 mL) were added to a 1-dram vial. The solution was sparged with $N_2$, followed by $H_2$ for 10 minutes each. The reduction was carried out under 1 atm $H_2$ at 60 °C overnight. Subsequently, the product was purified by silica flash chromatography to give 15 mg of a colorless oil (15% yield). Enantiomeric excess was determined by chiral GC to be 95.7%. LCMS: $[M+H]^+$ = 231. $^1H$ NMR (300 MHz, $CDCl_3$) δ 7.28, 4.06, 4.05, 4.05, 4.04, 4.02, 4.01, 4.00, 3.74, 2.87, 2.58, 2.57, 2.52, 2.51, 2.47, 2.44, 2.42, 2.39, 1.57, 1.54, 1.51, 1.50, 1.47, 1.45, 1.43, 1.33, 1.30, 1.28, 1.27, 0.92, 0.91, 0.90, 0.88. $^{13}C$ NMR (75 MHz, $CDCl_3$) δ 173.6, 77.4, 77.0, 76.6, 68.0, 51.8, 41.1, 36.5, 31.9, 29.57, 29.55, 29.5, 29.3, 25.5, 22.7, 14.1.

**Derivatization of natural 3*R*-OH-5*Z*-$C_{12:1}$-HSL.** First, the acyl chain of 3*R*-OH-5*Z*-$C_{12:1}$-HSL **(3)** was reduced with Pd/C to yield 3-OH-$C_{12}$-HSL **(4)**. 3*R*-OH-5*Z*-$C_{12:1}$-HSL (**3**; 0.2 mg) was dissolved in absolute ethanol (0.1 mL) and sparged with $N_2$ for 10 minutes. Simultaneously, 5% (m/v) of palladium on carbon (Aldrich, 205699) in absolute ethanol (0.1 mL) was sparged with $N_2$, followed by sparging with $H_2$. The solutions were combined and left to react at room temperature for 4 hours, after which time the solvent was removed under a stream of $N_2$. Using a protocol adapted from Thiel *et al.*,[8] the resulting **4** was filtered through cotton, then underwent methanolysis, 2% V/V $H_2SO_4$ in absolute methanol, at 60 °C overnight to yield methyl 3-hydroxydodecanoate **(5)**.[11] Excess acid was neutralized with sodium bicarbonate, then the solution was filtered through cotton in preparation for GC analysis.

**Chiral gas chromatography analysis.** To determine the absolute stereochemistry of **5**, chiral GC analysis was performed. Retention times of **2**, racemic 3-hydroxydodecanoic acid methyl ester (TRC, H939600), and **5** were compared. Separation of the *R*/*S* methyl ester enantiomers was performed on an Agilent 6890 GC fitted with an Agilent HP-Chiral column (30m, 0.32mm i.d., Agilent) and a flame ionization detector. 1 µL of each sample was injected in split-injection mode (10:1). The instrument was run at 115 °C isocratic for 90 minutes, followed by an increase to 220 °C (5 °C/min). $H_2$ carrier gas (4.0 mL/min) was used.

**$MmaR_{DSM5686}$ reporter assay.** An overnight culture of AWP370 reporter strain (AWP348+pAWP492) was subcultured to an optical density of 0.05 in fresh AMS containing kanamycin (50 ug/mL), 0.01% (w/v) yeast extract and 50mM MeOH in a 50 mL sterile conical tube. 0.5 mL of this culture was added to each of the wells of a 96-well deep well plate containing 5 µL of the appropriate acyl-HSL dissolved in acidified ethyl acetate. The plate was then incubated at 30 °C for 24 hours with shaking (200 rpm). 100 µL of each well was transferred to a black with clear flat bottom 96-well plate (Corning 3631) and red fluorescence was quantified at 570 nm excitation and 605 nm emission using a SpectraMax i3x plate reader. Absorbance at 600 nm was also quantified for normalization. Each condition was performed in triplicate. Results were analyzed using Graphpad Prism version 8.0.2.

### SUPPLEMENTARY FIGURES

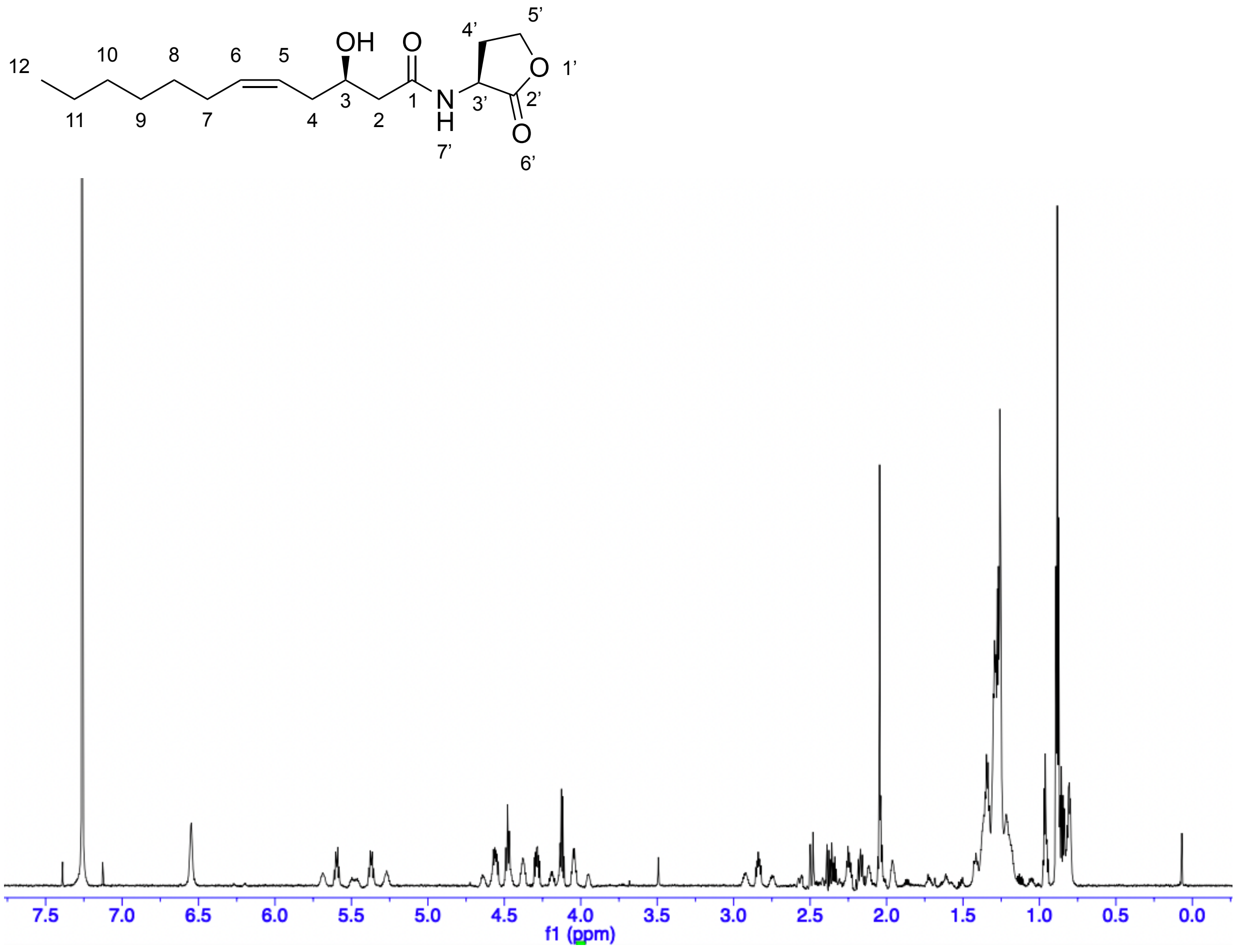

**Figure S1.** $^{1}H$ NMR spectrum of *3R*-OH-*5Z*-$C_{12:1}$-HSL in $CDCl_3$ (800 MHz).

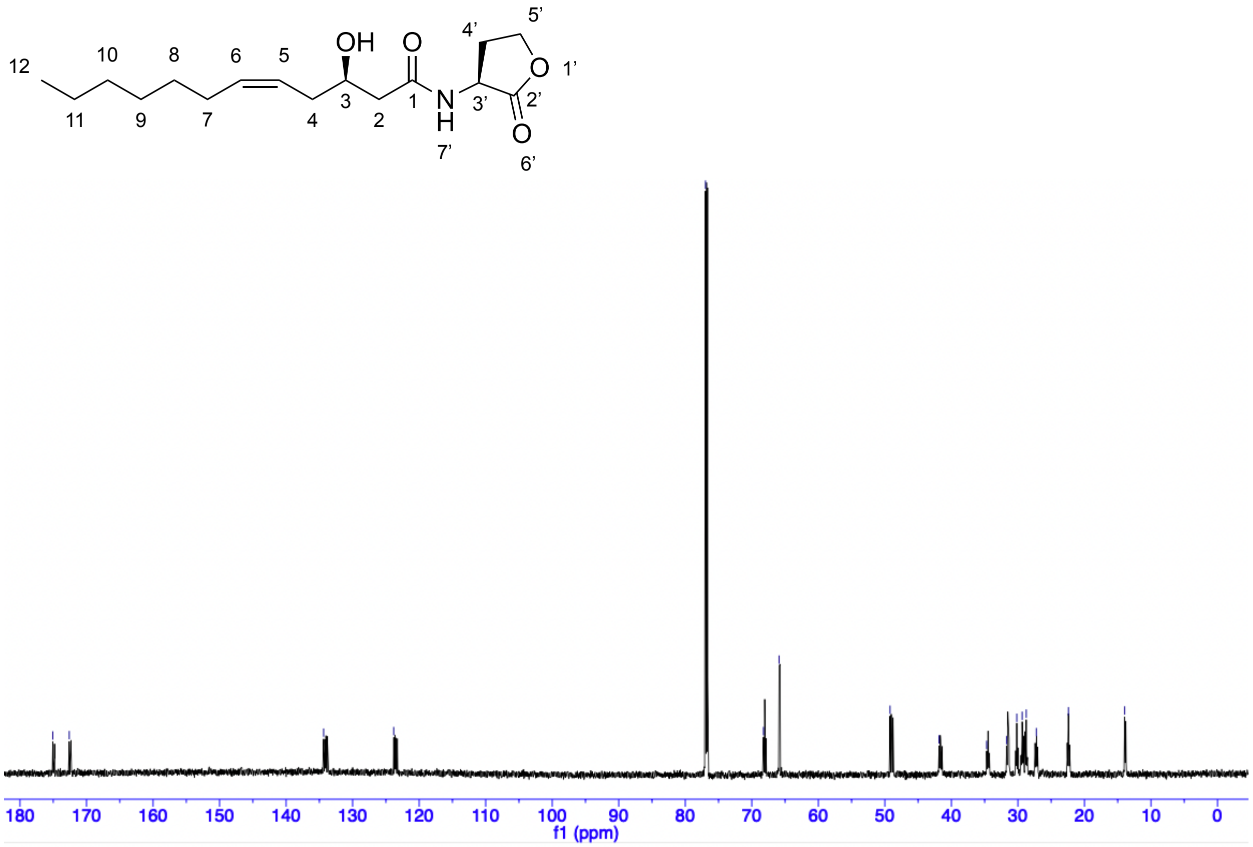

**Figure S2.** $^{13}$C NMR spectrum of *3R*-OH-*5Z*-$C_{12:1}$-HSL in $CDCl_3$ (200 MHz).

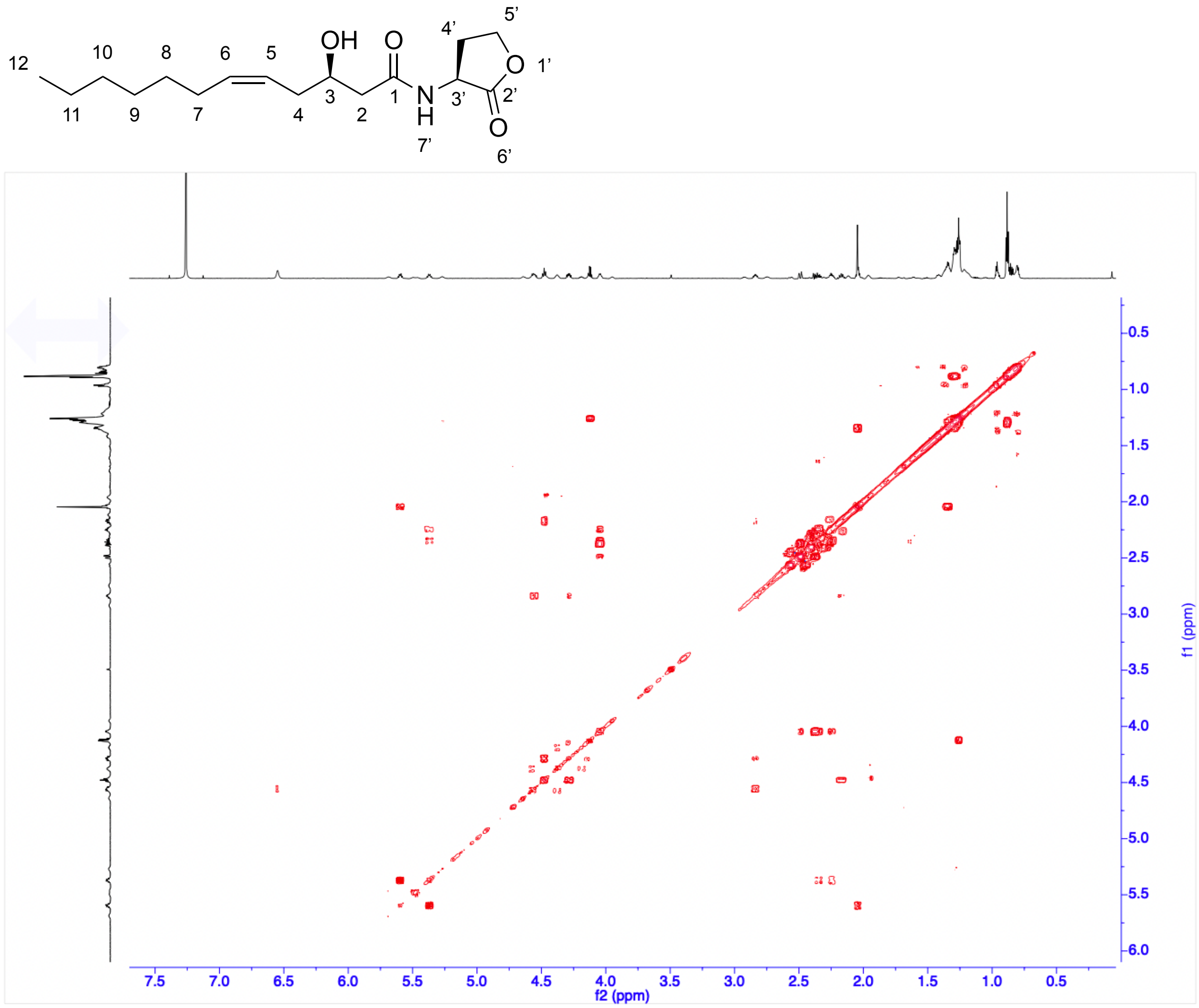

**Figure S3.** COSY spectrum of *3R*-OH-*5Z*-$C_{12:1}$-HSL in $CDCl_3$ (800 MHz).

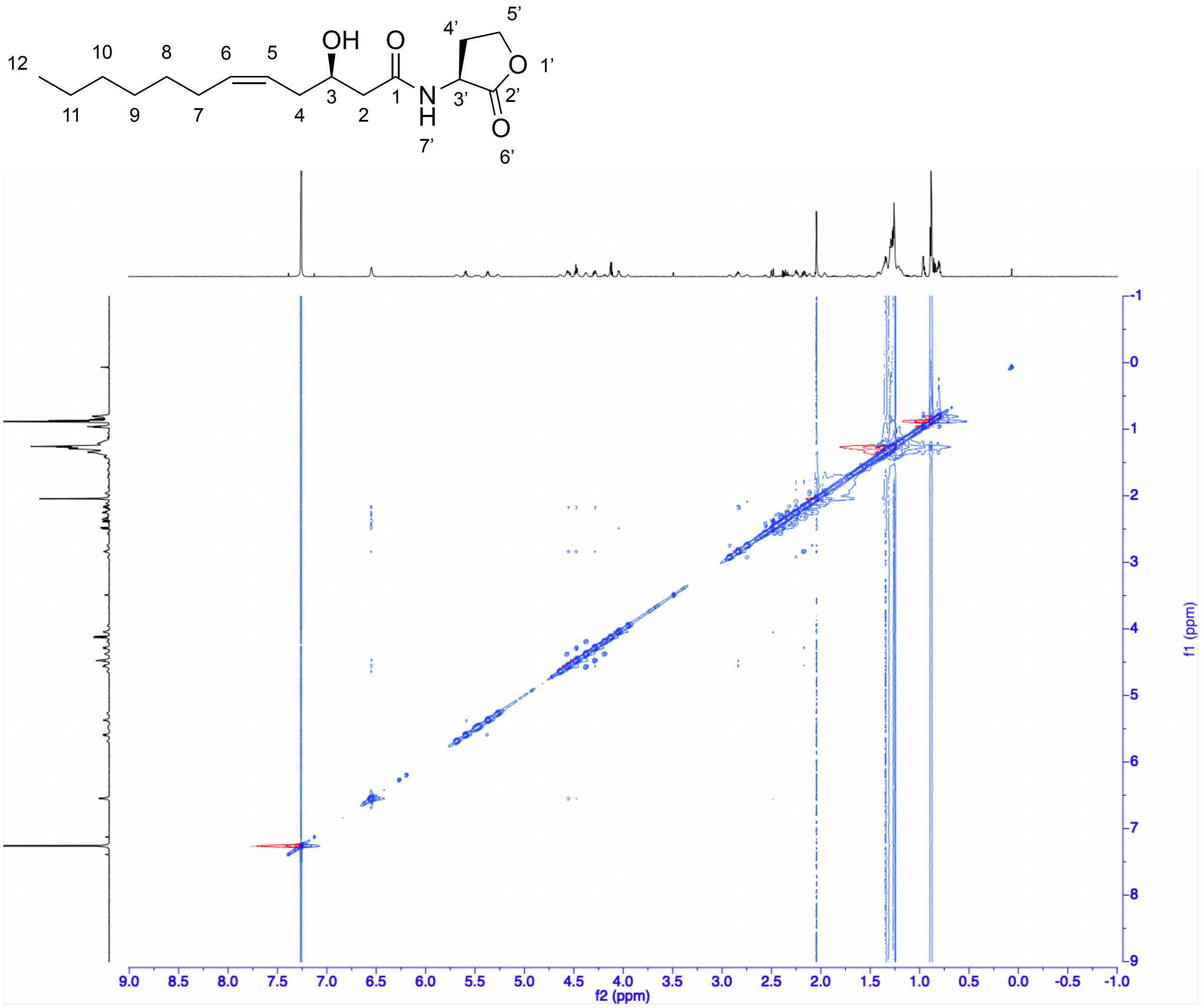

**Figure S4.** NOESY spectrum of 3*R*-OH-5*Z*-$C_{12:1}$-HSL in $CDCl_3$ (800 MHz).

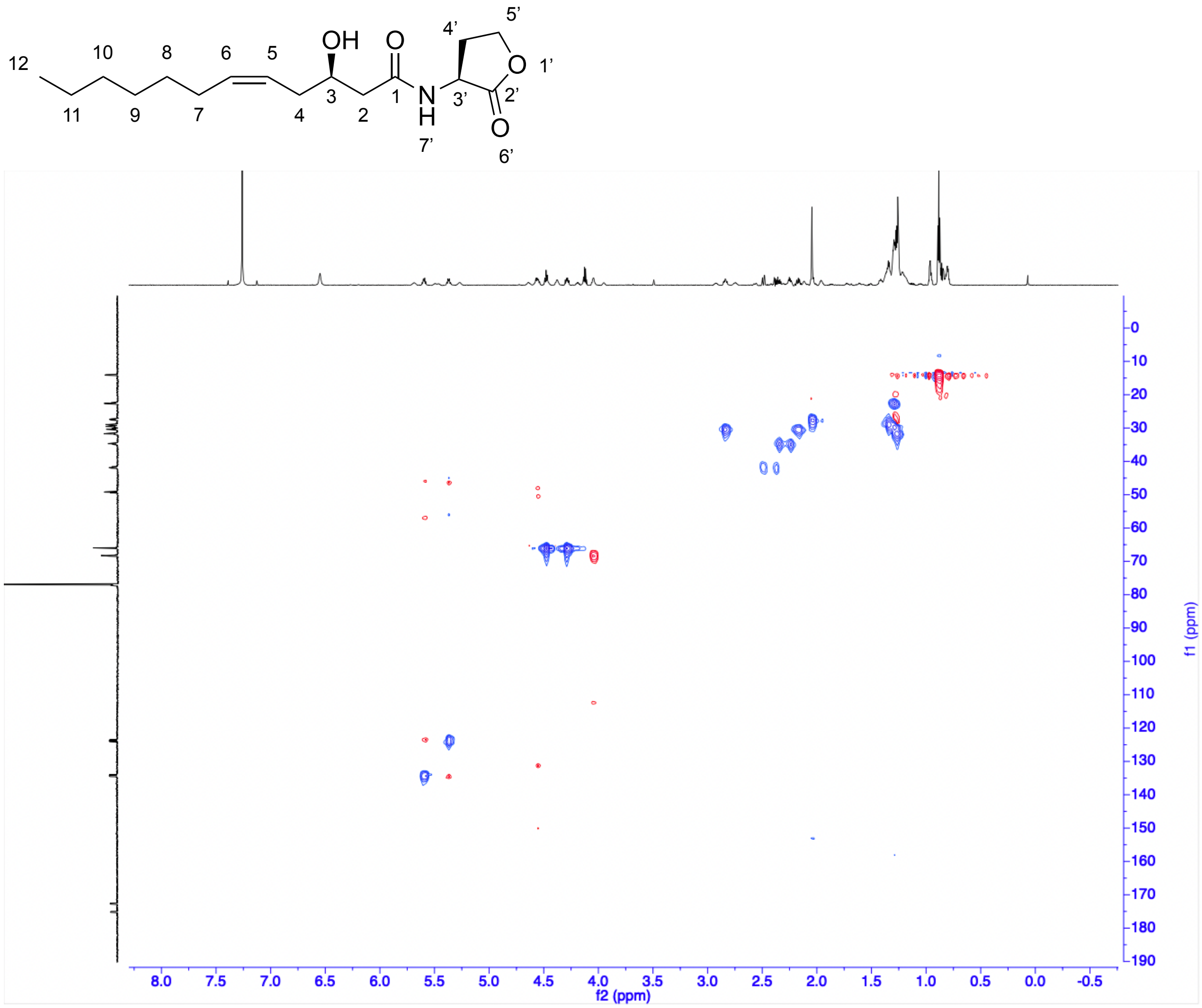

**Figure S5.** gHSQC spectrum of 3*R*-OH-5*Z*-$C_{12:1}$-HSL in $CDCl_3$ (800 MHz).

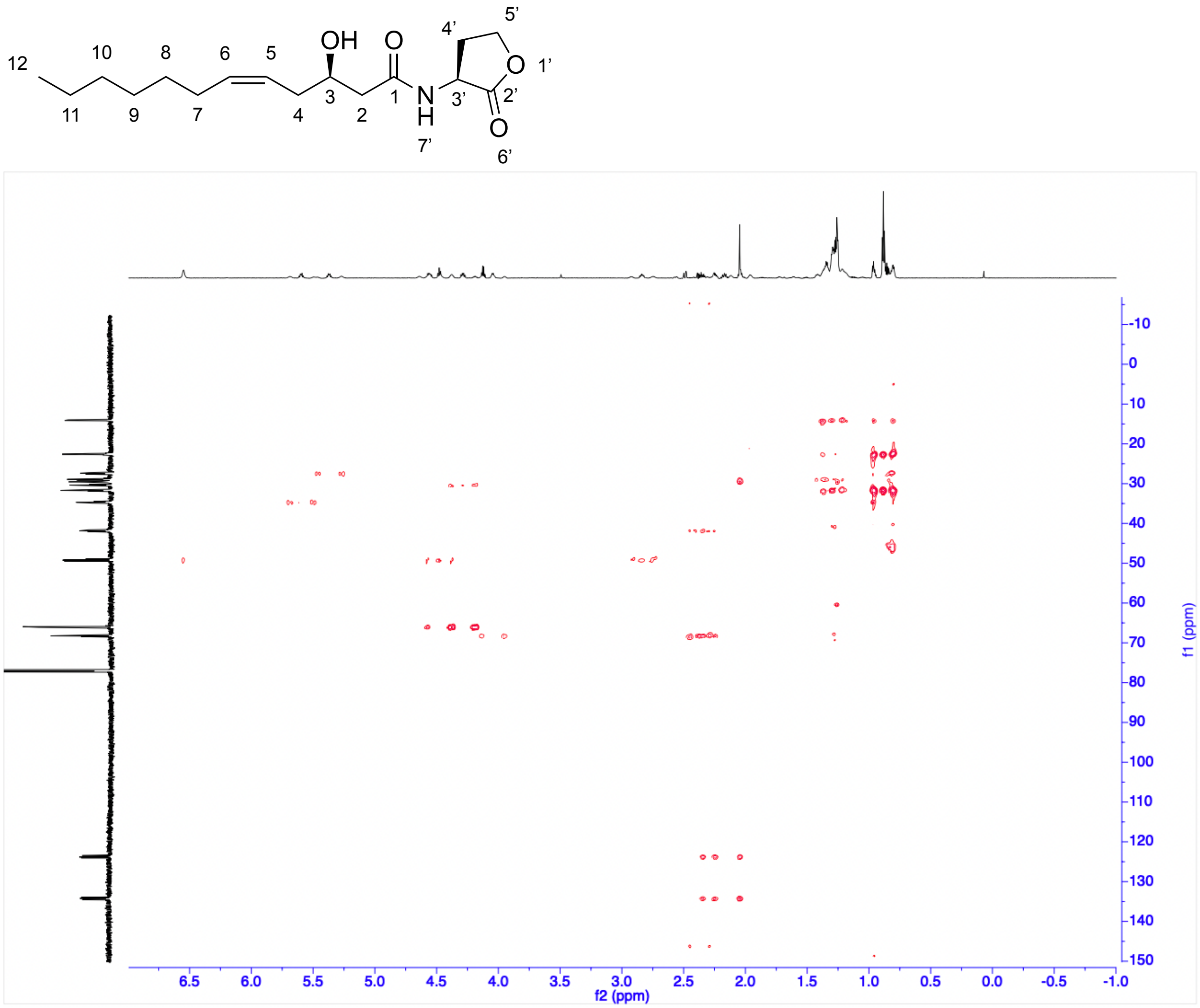

**Figure S6.** gHMBCAD spectrum of *3R*-OH-5*Z*-C$_{12:1}$-HSL in $CDCl_3$ (800 MHz).

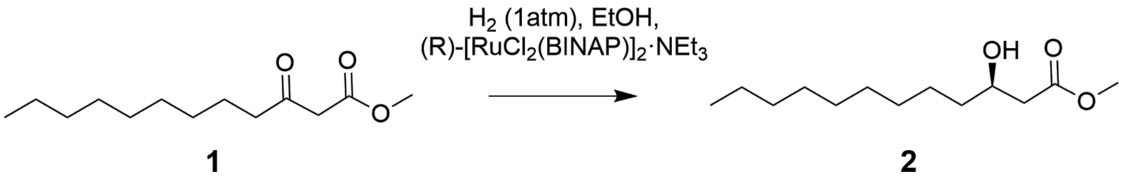

**Figure S7.** Synthesis of methyl 3*R*-OH-dodecanoate standard.

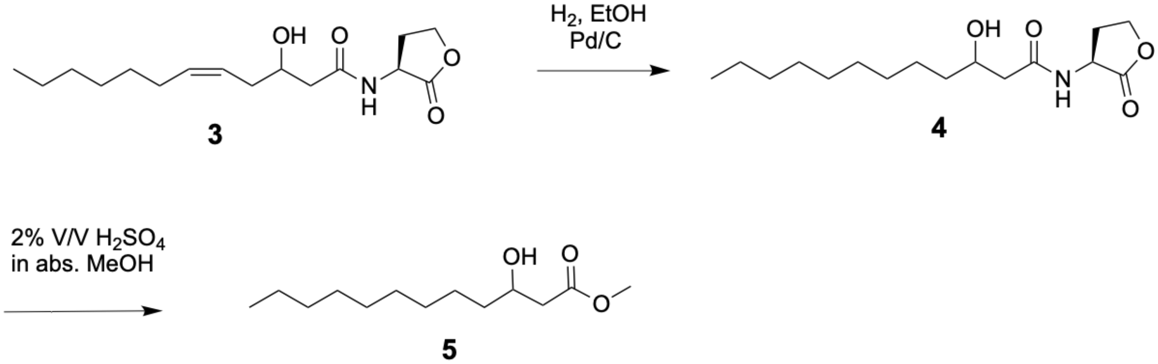

**Figure S8.** Derivatization of 3*R*-OH-5*Z*-$C_{12:1}$-HSL for chiral GC analysis.

A

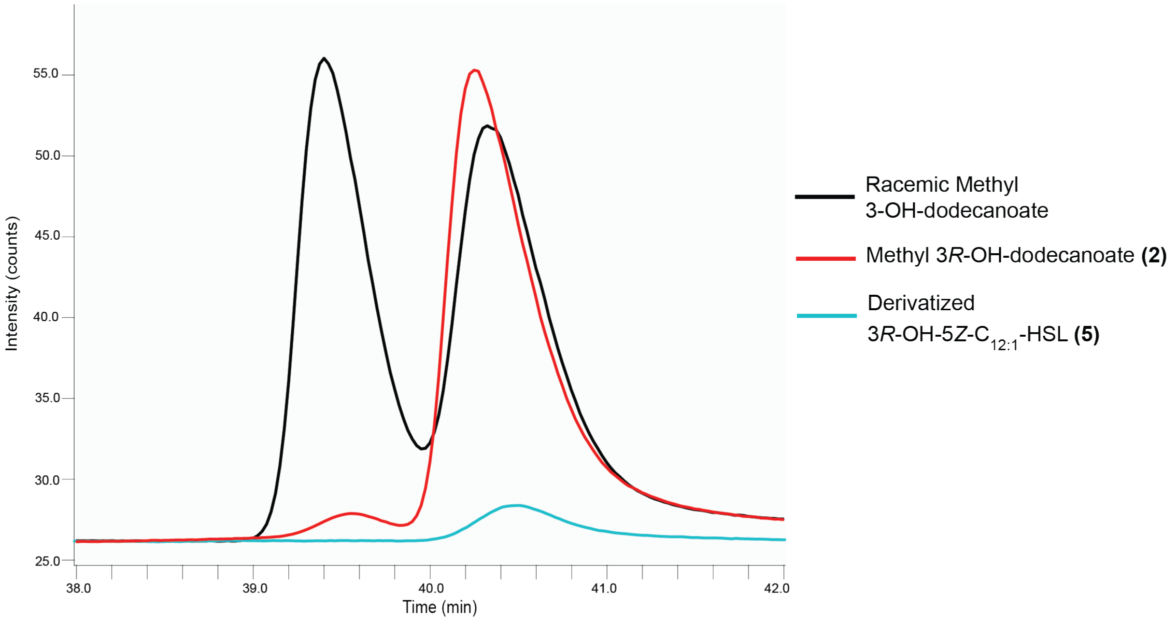

B

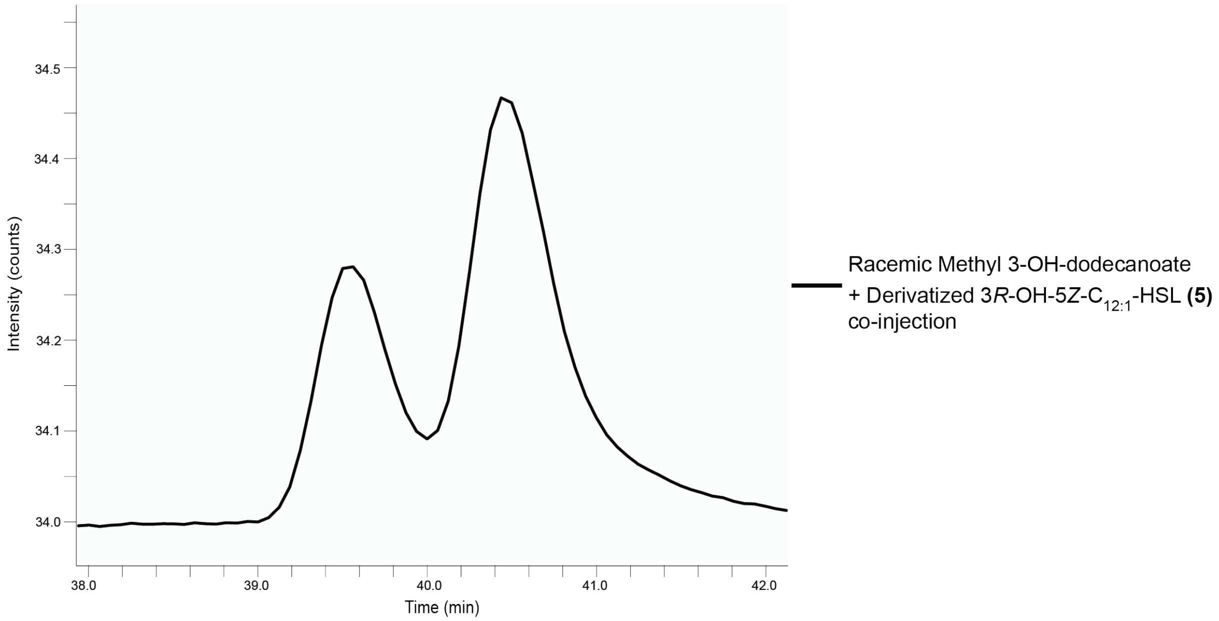

**Figure S9.** Gas chromatography analysis of derivatized natural 3*R*-OH-5*Z*-$C_{12:1}$-HSL, supporting the determined *R* stereochemistry of the 3-hydroxyl group. (A) GC trace overlays of racemic methyl 3-OH-dodecanoate, methyl 3*R*-OH-dodecanoate and derivatized natural 3*R*-OH-5*Z*-$C_{12:1}$-HSL. (B) Co-injection of racemic methyl 3-OH-dodecanoate and derivatized natural 3*R*-OH-5*Z*-$C_{12:1}$-HSL.

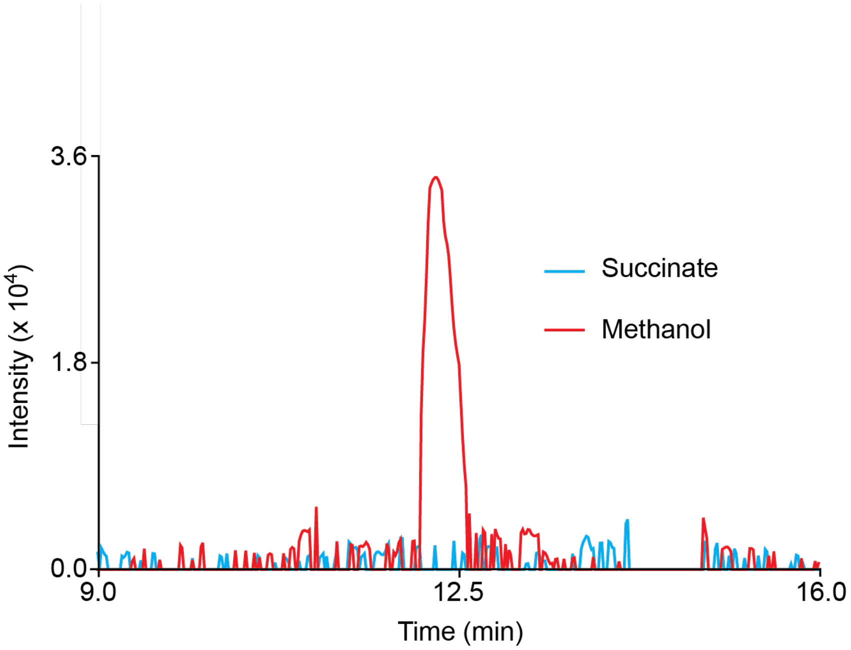

**Figure S10.** Extracted ion chromatogram of supernatant extracts of DSM5686 grown on the indicated carbon sources for *m/z* 298.2, corresponding to protonated 3*R*-OH-5*Z*-$C_{12:1}$-HSL. Mass tolerance ± 0.5 *m/z*.

### SUPPLEMENTARY TABLES

**Table S1.** MS/MS peak list for *m/z* 298 feature compared to commercial 3-oxo-$C_{12}$-HSL standard. The 15 most intense signals are shown for each sample.

| 3-oxo-C12-HSL | | 88A | | JCM2831 | | DSM 5686 | | AWP314 | |
|---|---|---|---|---|---|---|---|---|---|
| *m/z* | Intensity | *m/z* | Intensity | *m/z* | Intensity | *m/z* | Intensity | *m/z* | Intensity |
| 302.3053 | 1.95E+05 | 102.0564 | 4.02E+05 | 102.0566 | 1.99E+05 | 102.0569 | 1.33E+06 | 102.0565 | 1.51E+06 |
| 298.2013 | 5.53E+04 | 298.2016 | 3.22E+05 | 298.2019 | 1.50E+05 | 298.2021 | 1.08E+06 | 298.2018 | 1.24E+06 |
| 197.1533 | 4.60E+04 | 280.191 | 1.29E+05 | 280.1914 | 6.02E+04 | 280.1914 | 3.75E+05 | 280.191 | 4.13E+05 |
| 102.0565 | 4.01E+04 | 179.1432 | 6.63E+04 | 95.0876 | 3.19E+04 | 95.0878 | 2.38E+05 | 95.0875 | 2.76E+05 |
| 299.205 | 1.03E+04 | 299.2048 | 6.45E+04 | 299.2049 | 3.10E+04 | 179.1434 | 2.02E+05 | 299.2049 | 2.36E+05 |
| 155.1429 | 8.69E+03 | 155.1432 | 5.43E+04 | 179.1433 | 3.10E+04 | 137.1332 | 1.94E+05 | 137.1328 | 2.32E+05 |
| 198.157 | 5.76E+03 | 95.0873 | 5.18E+04 | 161.1327 | 2.46E+04 | 155.1436 | 1.83E+05 | 179.1432 | 2.27E+05 |
| 284.2951 | 5.42E+03 | 137.1328 | 5.03E+04 | 137.1331 | 2.46E+04 | 161.1329 | 1.60E+05 | 155.1432 | 2.13E+05 |
| 240.196 | 4.44E+03 | 161.1325 | 4.83E+04 | 155.1435 | 2.45E+04 | 81.0729 | 1.41E+05 | 161.1326 | 1.76E+05 |
| 270.2073 | 3.73E+03 | 81.0724 | 3.05E+04 | 81.0727 | 1.69E+04 | 281.1947 | 7.23E+04 | 81.0725 | 1.69E+05 |
| 98.0614 | 2.93E+03 | 281.1944 | 2.47E+04 | 302.3058 | 1.34E+04 | 103.06 | 7.12E+04 | 281.1943 | 8.05E+04 |
| 302.2688 | 2.41E+03 | 103.0596 | 2.21E+04 | 281.1949 | 1.25E+04 | 119.0868 | 6.19E+04 | 103.0597 | 7.98E+04 |
| 74.0632 | 2.29E+03 | 252.1961 | 1.63E+04 | 103.0597 | 1.04E+04 | 252.1963 | 5.31E+04 | 119.0862 | 6.96E+04 |
| 280.1902 | 2.22E+03 | 234.1857 | 1.62E+04 | 119.0865 | 9.68E+03 | 109.1026 | 5.24E+04 | 109.1024 | 5.95E+04 |
| 252.1969 | 2.17E+03 | 119.0863 | 1.48E+04 | 74.0632 | 8.12E+03 | 74.0635 | 5.17E+04 | 74.0633 | 5.61E+04 |

**Table S2.** NMR assignments for *3R*-OH-*5Z*-$C_{12:1}$-HSL in $CDCl_3$

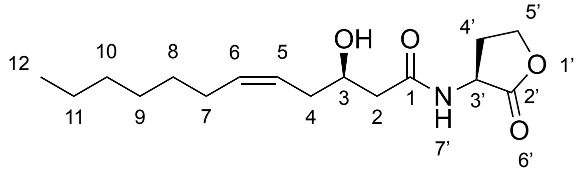

| **Position** | **$\delta_c$, type** | **$\delta_H$ (*J* in Hz)** | **HMBC** |
|---|---|---|---|
| 5' | 65.9, $CH_2$ | 4.28, 4.48, m | 2', 3', 4' |
| 4' | 29.4, $CH_2$ | 2.84, m | 2', 3', 5' |
| 3' | 49.2, CH | 4.56, dd (6.7) (1.7) | 2', 4', 5' |
| 2' | 179.3, C | | 3', 4', 5' |
| 1 | 176.5, C | | 2, 3 |
| 2 | 41.9 $CH_2$ | 2.38, 2.50, dd (5.8) | 1, 3 |
| 3 | 68.3, CH | 4.04, tt (7.7) | 1, 2, 4 |
| 4 | 34.7, $CH_2$ | 2.25, 2.36, dd (6.9) | 2, 5, 6 |
| 5 | 127.8, CH | 5.59, (8.9) | 4, 6, 7 |
| 6 | 138.2, CH | 5.36, (8.9) | 4, 5, 7, 8 |
| 7 | 27.2, $CH_2$ | 2.17, q (8.9) | 5, 6, 8, 9 |
| 8 | 28.8, $CH_2$ | 2.05, tt (7.2) | 6, 7, 9, 10, 11 |
| 9 | 31.7, $CH_2$ | 1.34, m | 7, 8, 10, 11, 12 |
| 10 | 30.2, $CH_2$ | 1.28, m | 8, 9, 11 |
| 11 | 22.4, $CH_2$ | 1.26, m | 8, 9, 10, 12 |
| 12 | 14.0, $CH_3$ | 0.88, t (7.0) | 9, 10, 11 |
| NH (7') | | 6.55 | |

**Table S3.** Marfey's analysis of HSL portion of *3R*-OH-*5Z*-$C_{12:1}$-HSL.

| **Sample** | **Retention time (min)** |
|---|---|
| Marfey's derivatized *D*-HSL standard (*m/z* 372) | 13.6 |
| Marfey's derivatized *L*-HSL standard (*m/z* 372) | 15.2 |
| Marfey's derivatized *3R*-OH-*5Z*-$C_{12:1}$-HSL hydrolysate (*m/z* 372) | 15.4 |

**Table S4.** Strains used in this study.

| Strain | Puri Lab Strain Collection Number | Description[a] | Reference |
|---|---|---|---|
| *E. coli* TOP10 | EAWP2 | F– *mcrA* Δ(*mrr-hsdRMS-mcrBC*) Φ80*lacZ*ΔM15 Δ*lacX74 recA1 araD139* Δ(*ara leu*) 7697 *galU galK rpsL* ($Str^R$) *endA1 nupG* | Invitrogen |
| *E. coli* S17-1 λpir | EAWP3 | Donor strain. $Tp^R$ $Sm^R$ *recA thi pro hsd*($r^- m^+$) RP4-2-Tc::Mu::Km Tn7 λ*pir* | [7] |
| *Methylobacterium* sp. strain 88A | AWP105 | Pink pigmented facultative methylotroph. | [9] |
| *M. radiotolerans* JCM2831 | AWP120 | Pink pigmented facultative methylotroph. Source of $mmaI_{JCM2831}$ gene. | [10] |
| *M. extorquens* AWP227 | AWP227 | Heterologous expression strain. Derivative of *M. extorquens* PA1. Δ(*celABC*-Mext_1370) Δ*mlaRI* | [2] |
| *M. fujisawaense* DSM5686 | AWP269 | Pink pigmented facultative methylotroph. | [11] |
| *M. extorquens* AWP227 + pAWP417 | AWP314 | Heterologous expression of $mmaI_{JCM2831}$ | This study |
| *M. extorquens* AWP348 | AWP348 | Insertion mutant containing $mmaR_{DSM5686}$ downstream of the *mlaR* promoter. Δ(*celABC*-Mext_1370) Δ*mlaRI::*$mmaR_{DSM5686}$ | This study |
| *M. extorquens* AWP348 + pAWP492 | AWP370 | $MmaR_{DSM5686}$ reporter strain. | This study |
| *M. fujisawaense* AWP374 | AWP374 | Δ*mmaI* | This study |

[a]IMG/M gene locus tags: Δ$mlaRI_{PA1}$, Mext_4513-4; $mmaR_{DSM5686}$, Ga0373205_3344; $mmaI_{DSM5686}$, Ga0373205_3345; $mmaI_{JCM2831}$: Mrad2831_5763.

**Table S5.** Plasmids used in this study.

| Plasmid | Puri Lab Plasmid Collection Number | Description[a] | Reference |
|---|---|---|---|
| pAWP78 | pAWP78 | IncP-based expression vector. | [12] |
| pAWP227 | pAWP227 | pCM433kanT containing flanks to knock out *mlaRI* in PA1. | [2] |
| pAWP274 | pAWP274 | Inserting *mmaR*$_{DSM5686}$ into AWP227. | This study. |
| pCM433kanT | pAWP285 | Sucrose counterselection vector for creating unmarked deletion mutants. | [12] |
| pMRE-Tn7-135 | pAWP398 | Source of *mScarlet* gene. | [13] |
| pAWP417 | pAWP417 | Expressing *mmaI*$_{JCM2831}$ gene under the *dnaG*$_{PA1}$ promoter (400 bp upstream sequence). | This study |
| pAWP429 | pAWP429 | pCM433kanT containing flanks to knock out the *mmaI* gene in DSM5686. | This study |
| pAWP492 | pAWP492 | Expressing mScarlet under the *mmaI*$_{DSM5686}$ promoter (400 bp upstream sequence). | This study |

[a]IMG/M gene locus tags: Δ*mlaRI*$_{PA1}$, Mext_4513-4; *mmaR*$_{DSM5686}$, Ga0373205_3344; *mmaI*$_{DSM5686}$, Ga0373205_3345; *mmaI*$_{JCM2831}$, Mrad2831_5763; *dnaG*$_{PA1}$, Mext_0611.

**Table S6.** Primers used in this study. Homology regions used for Gibson Assembly are bolded.

| Primer Name | Sequence (5' to 3') | Description[a] |
|---|---|---|
| oAWP186_ 433KTV1_fwd | ATGTGCAGGTTGTCGGTGTC | For amplifying the pCM433kanT backbone. oAWP186 and 160 were used to amplify one piece, and oAWP159 and 187 were used to amplify the other. |
| oAWP160_ 433KTV1_rev | **ATAAAGGTGAATCCCATAGGGCAGGAGCTA**TAATCTCGAGTCCCGTCAAG | |
| oAWP159_433KTV2_fwd | TAGCTCCTGCCCTATGGGAT | |
| oAWP187_433KTV2_rev | TGGTAACTGTCAGACCAAGTTTACTC | |
| oAWP259_78V_fwd1 | TTGTCGGGAAGATGCGTGAT | For amplifying the pAWP78 backbone. |
| oAWP254_78V_rev1 | CAGCTCACTCAAAGGCGGTA | |
| oAWP994_PdnaG_fwd | **AGCGCGTACTCCGTCCCCCGAACGTTGCAT**GGGGACTCTGCTGGAAGCGG | For amplifying the *dnaG*$_{PA1}$ promoter (400 bp upstream sequence) for heterologous expression of *mmaI*$_{JCM2831}$. |
| oAWP995_PdnaG_rev | **TCCACGCGATCCGCTTCCAGCAGAGTCCCC**ATGATCCATGTCGTGACTGC | |
| oAWP1055_fwd | CATGGCCCGGCAAAATCATC | For opening pAWP227 between the up and down flanks to insert *mmaR*$_{DSM5686}$. oAWP1055 was used with oAWP160 and oAWP1056 was used with oAWP159. |
| oAWP1056_rev | CGGACGCGGCGACGGGTGCG | |
| oAWP1060_274I_fwd | **ATGAGAGTGCGATGATTTTGCCGGGCCATG**CCGCATTCGAAGCACCTGGA | For amplifying the *mmaR*$_{DSM5686}$ to insert between the up and down flanks of pAWP227. |
| oAWP1061_274I_rev | **ACGGATGCGGCGCACCCGTCGCCGCGTCCG**CTACGTGATCAGGCCGGCAC | |
| oAWP1086_417I_fwd | **GAAGGATCAGATCACGCATCTTCCCGACAA**ATGATCCATATCGTCACACCCGCC | For amplifying *mmaI*$_{JCM2831}$. |
| oAWP1087_417I_rev | **TCCACGCGATCCGCTTCCAGCAGAGTCCCC**TCAGGCCACCAGGTAAGCGGGTT | |
| oAWP1448_429U_fwd1 | **TTTTGCCGGGCCATG**TTCAAACGGCAGAGCTTTCC | For amplifying flanks to knock out *mmaI* in DSM5686. |
| oAWP1449_429U_rev1 | **GGCGAGCCGGGCGAA**CACGAAGTAACCTCACGTTG | |

| | | |
|---|---|---|
| oAWP1450_429D_fwd1 | **CGTGCATCACGACACCGACAACCTGCACAT**AACTCGGTCCTGGTCTCCGA | |
| oAWP1446_429D_rev1 | **ATAACCGTATTACCGCCTTTGAGTGAGCTG**AGTACCCGGTCTACCTCTCG | |
| oAWP1535_450I2_fwd | ATGGTGAGCAAGGGCGAGG | For amplifying *mScarlet* from pMRE-Tn7-135. |
| oAWP1504_450I2_rev | **GAAGGATCAGATCACGCATCTTCCCGACAA**CTTGTACAGCTCGTCCATGCC | |
| oAWP1617 fwd 5686 upstream I gene | **ATAACCGTATTACCGCCTTTGAGTGAGCTG**CAACTCCTGCTGATGAACAGC | For amplifying the *mmaI*$_{DSM5686}$ promoter (400 bp upstream sequence). |
| oAWP1618 rev 5686 upstream I gene | **ATAACCGTATTACCGCCTTTGAGTGAGCTG**CAACTCCTGCTGATGAACAGC | |

[a]IMG/M gene locus tags: *mmaR*$_{DSM5686}$, Ga0373205_3344; *mmaI*$_{DSM5686}$, Ga0373205_3345; *mmaI*$_{JCM2831}$, Mrad2831_5763; *dnaG*$_{PA1}$, Mext_0611.
